# Using Bayesian inference to characterize cell-size regulation across organisms

**DOI:** 10.64898/2026.10.14.759273

**Authors:** Kuheli Biswas, Michael Sheinman, Ariel Amir

## Abstract

How organisms regulate cell size and whether this varies with growth conditions remains a fundamental question. Inferring how cells regulate their size requires separating biological control from statistical effects. Yet regulatory strategies are commonly classified by estimating the size-control parameter, *α*, from single-cell measurements using linear regression. We show that this conventional estimator is systematically biased by measurement noise, lineage-to-lineage extrinsic variability, and finite lineage length. Consequently, the same underlying regulatory mechanism can be misclassified as sizer-, adder-, or timer-like. We develop a Bayesian framework that jointly infers *α* and the amplitudes of intrinsic, extrinsic, and measurement noise while accounting for lineage heterogeneity and finite sampling. Across simulations spanning experimentally relevant noise levels and lineage lengths, the framework recovers *α* and outperforms conventional regression, particularly for short and heterogeneous lineages. Reanalysis of diverse bacterial and eukaryotic datasets collected under multiple growth conditions indicates that statistical bias accounts for much of the reported variation in *α* across organisms and growth conditions. After accounting for these biases, the datasets are broadly consistent with adder-like regulation. Thus, our results provide a general framework for distinguishing genuine biological differences from noise-induced changes in inferred regulatory strategy and revise how cell-size regulation should be inferred from lineage data.

## 2 Introduction

Cell-size homeostasis is a fundamental requirement for cellular proliferation, yet the principles by which cells regulate their size remain unresolved. Although cell-size control is implemented through diverse molecular pathways [1, 2], evolutionarily distant organisms—including bacteria, archaea, and eukaryotes—may nevertheless share a common phenomenological (coarse-grained) description of cell-size regulation [3, 4, 5, 6, 7]. To enable quantitative comparisons of cell-size regulation across organisms, several seminal studies introduced phenomenological models that classify cell-size regulation according to how division size depends on birth size [8, 9, 10, 11]. In a commonly used small noise approximation, the division size 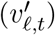 of a cell in lineage *ℓ* at generation *t* is related to its birth size (*v*_*ℓ,t*_) by

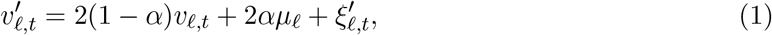

where *α* is the cell-size regulation parameter, *µ*_*ℓ*_ is the mean birth size of lineage *ℓ*, and 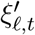 is zeromean Gaussian noise. It follows from Eq. (1) that stable size homeostasis requires (0 < *α* < 2). This range includes several canonical regulatory strategies: *α* = 1 corresponds to a *sizer*, for which cells divide upon reaching a characteristic size; *α* = 0.5 corresponds to an *adder*, for which cells add a characteristic size between birth and division; and *α* → 0^+^ corresponds to timer-like regulation, for which exponentially growing cells divide after an approximately characteristic time. In singlecell studies, *α* is typically inferred by pooling cell-size data across lineages and performing a linear regression of division size on birth size; we refer to this method as the *pooled estimator*.

The application of the pooled estimator has yielded inconsistent estimates of cell-size regulation among datasets from the same organism. In *E. coli*, pooled estimates of *α* span approximately 0.20– 0.9, encompassing timer-, adder-, and sizer-like regulation. Estimates are generally adder-like under fast growth [12, 13, 3, 14], whereas the data of Kennard *et al*. yield timer-like values [15]. Under slow growth, reported estimates range from adder-like [3] to increasingly sizer-like [16, 14, 17]. Similar variability is observed in the fission yeast *S. pombe*, for which pooled estimates range from approximately 0.56 to 0.8 across growth conditions, indicating regulation intermediate between an adder and a sizer [18, 19, 20]. Taken together, these disparate, condition-dependent pooled estimates indicate that no consensus exists regarding the underlying cell-size regulation parameter, even within a single organism. Nevertheless, the apparent shift from adder-like regulation has motivated organism- and condition-specific biological explanations, including increased degradation of FtsZ, a key division protein, in slow-growing *E. coli* [14, 17, *21] and geometry-dependent control of mitotic entry through the Cdr2 pathway in fission yeast [2, 22]*.

*Before delving into biological explanations for deviations from adder regulation, we first examine the inference method of α* from Eq. (1) as statistical errors can produce apparent changes in regulatory strategy even when the underlying regulation remains unchanged. Single-cell data are subject to measurement noise and extrinsic noise, which generate lineage-to-lineage variability in average cell size. We show that both sources of variation systematically bias the commonly used pooled estimator (see Fig. 1).

**Figure 1:**
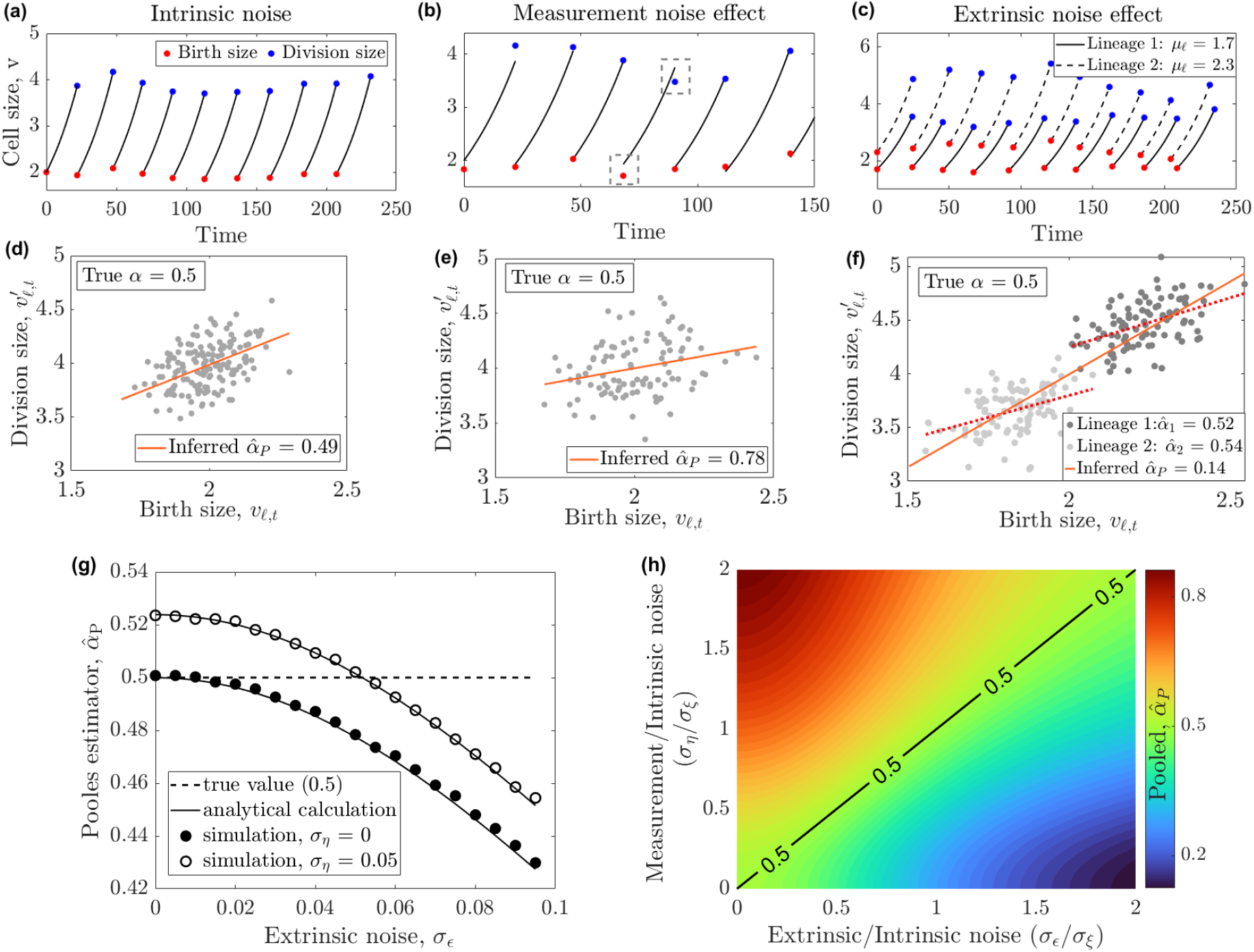
Pooled estimator is biased under measurement and extrinsic noise. (**a–c**) Representative cell-size trajectories generated with intrinsic noise alone (**a**), measurement noise (**b**), and extrinsic noise (**c**). Measured birth and division sizes are indicated in red and blue, respectively. Measurement noise results in deviation between true and measured sizes [gray dashed box in (**b**)]. Extrinsic noise produces lineage-to-lineage variability in average birth size [examples of two lineages are presented in solid and dotted lines in (**c**)]. (**d–f** ) Corresponding regressions of division size on birth size for intrinsic noise alone (**d**), measurement noise (**e**), and extrinsic noise (**f** ). Extrinsic noise in (**f** ) creates a discrepancy between the within-lineage regression slopes and the pooled regression slope, which is related to Simpson’s paradox [29]. Although the true regulation parameter is *α* = 0.5, measurement and extrinsic noise shift the pooled estimate away from its true value. (**g**) We validate the analytical prediction for the pooled estimator 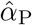 in Eq. 9 against simulations over a range of extrinsic and measurement noise levels. The dashed line denotes the true value, *α* = 0.5. (**h**) Heat map of 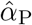 as a function of extrinsic and measurement noise, each normalized by the intrinsic noise, for the true value of *α* = 0.5. Measurement noise biases 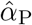 upward, whereas extrinsic noise biases it downward. The two biases cancel when 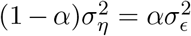, such that the predicted pooled estimate equals the true value. For *α* = 0.5, the condition reduces to *σ*_*η*_ = *σ*_*ϵ*_.

To overcome the bias in *α* from lineage-to-lineage variability, a recent study [23] proposed a different inference of *α* than the pooled estimator, where *α* is inferred separately for each lineage and then averaged across lineages, which we refer to as the *lineage-wise estimator*. However, we show that the lineage-wise estimator is biased upward for short lineages and that measurement noise introduces an additional upward bias. The predicted finite-length bias is also apparent in single-cell datasets from *E. coli* and mouse leukemia cells (see Fig. 2).

**Figure 2:**
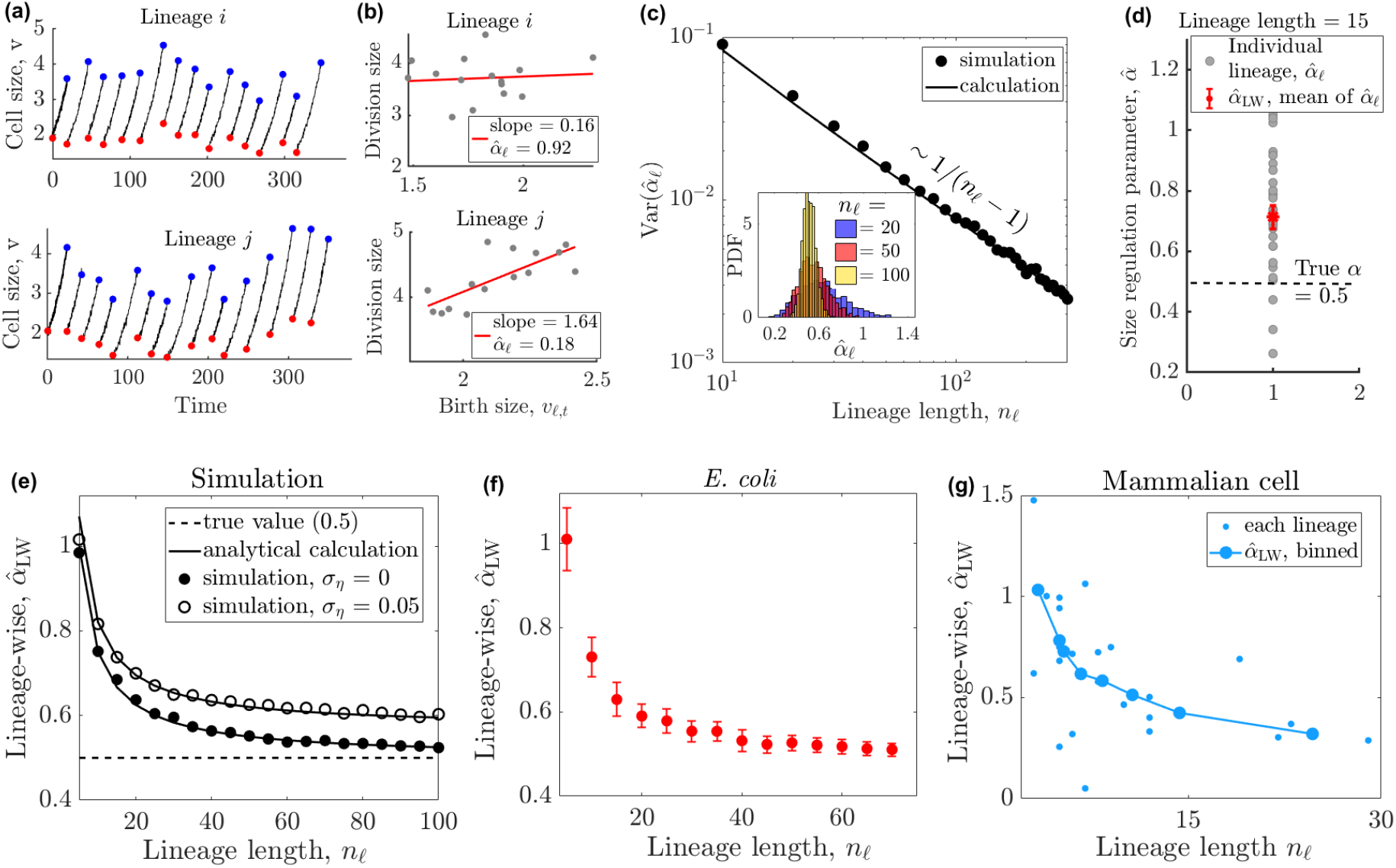
Lineage-wise estimator is biased under measurement noise and short lineages. (**a**) Representative cell-size trajectories from two lineages, *i* and *j*, with measurement noise (CV_*η*_ = 0.05) and lineage length *n*_*ℓ*_ = 15. Red and blue circles indicate birth and division sizes, respectively. (**b**) Corresponding linear regressions of division size on birth size for the lineages shown in (**a**) vary widely across lineages (see 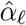 in legend). (**c**) The variance of 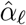 scales approximately as 1*/*(*n*_*ℓ*_ − 1). An analytical form of 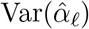 is provided in Appendix Eq. (B.55). In the inset, we show the distribution of 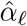 for three different values of *n*_*ℓ*_, which is narrower as *n*_*ℓ*_ increases. (**d**) 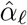 (gray circles) and 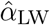 (red), mean of 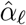 for lineage lengths *n*_*ℓ*_ = 15 overestimates the true value of *α* = 0.5 (dashed line). (**e**) Lineage-wise estimator 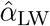 as a function of lineage length in simulations with and without measurement noise. The analytical prediction in Eq. (11) agrees with the simulation results and shows that short lineages bias the estimate upward, while measurement noise introduces an additional bias. The dashed line denotes the true value, *α* = 0.5. A similar lineage-length dependence is observed in experimental data from single-cell *E. coli* (**f** ) [32] and mouse leukemia cells (**g**) [23], consistent with the finite-lineage bias predicted by our model.

We develop a Bayesian inference framework for the cell-size model that explicitly accounts for intrinsic noise within lineages [Eq. (1)], extrinsic variability between lineages, and measurement noise. Within this framework, we jointly infer the size-regulation parameter *α*, the lineage-specific mean birth sizes, and the amplitudes of the different noise sources from the posterior distribution. Parameter estimates are reported as posterior means with 95% credible intervals. We benchmark the method using simulated data and show that it provides bias-corrected inference of *α*, particularly for short and heterogeneous lineages subject to measurement noise (Fig. 3).

**Figure 3:**
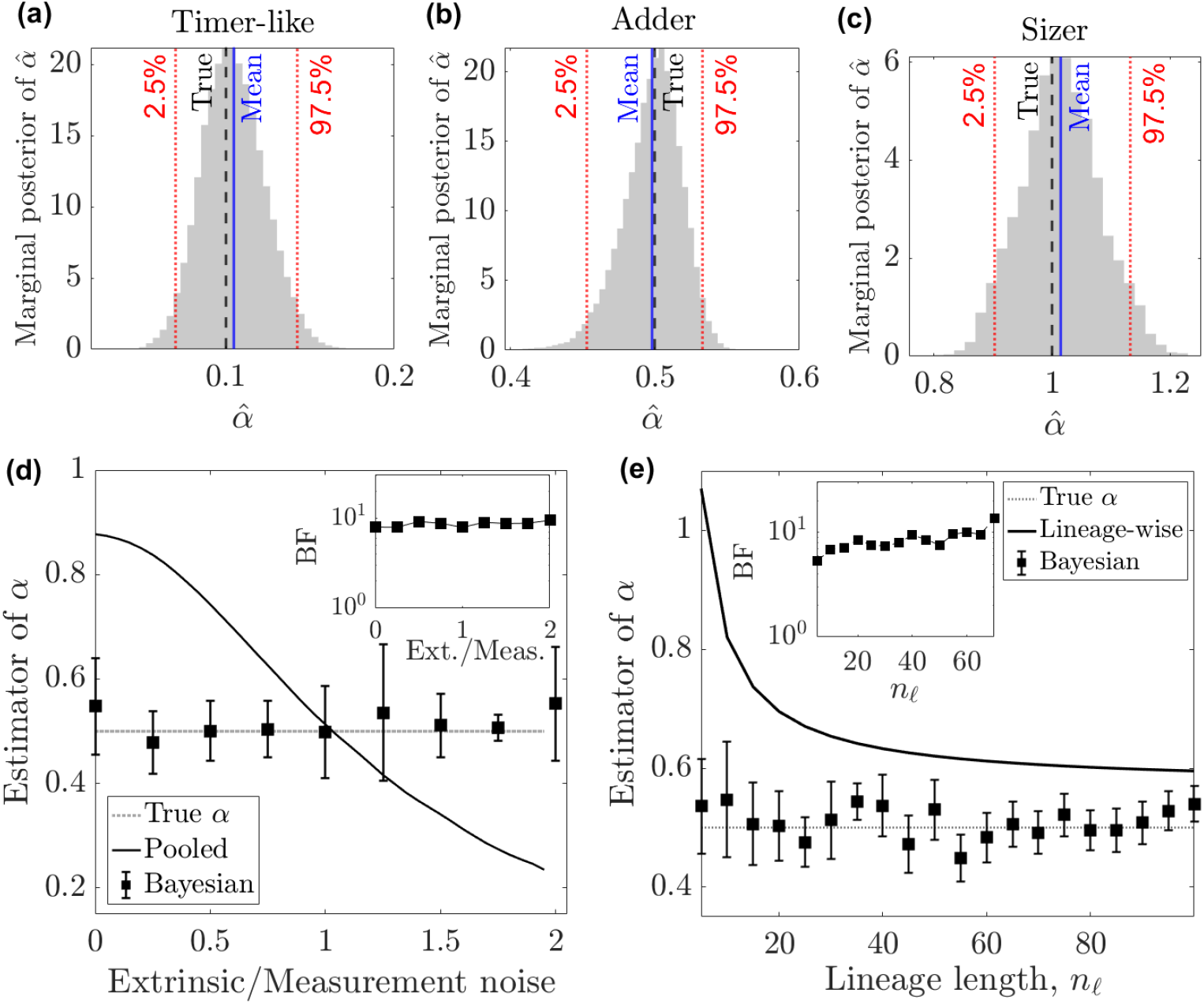
Bayesian inference provides a bias-corrected inference of *α*. (**a–c**) Representative marginal posterior distributions of the size-regulation parameter *α* for true values *α* = 0.1, 0.5, and 1.0, respectively. The solid blue line denotes the posterior mean, the dashed black line denotes the true value, and the red dotted lines mark 95% credible intervals, defined by the 2.5th and 97.5th posterior percentiles. (**d**) Estimated *α* as a function of the ratio between extrinsic and measurement. The conventional pooled estimator (solid line) deviates substantially from the true value, whereas the Bayesian estimate (squares) remains centered near *α* = 0.5. (**e**) Estimates of *α* as a function of lineage length. The conventional lineage-wise estimator (solid line) is biased upward, particularly for short lineages, whereas the Bayesian estimate remains close to the true value. The insets show the Bayes factor (BF) comparing the adder model (*α* = 0.5) against mixed strategies (*α* ≠ 0.5) at each noise ratio in **(d)** and lineage length in **(e)**. A BF of 10 indicates strong evidence for the adder model. For the simulations, we generated 50 lineages using the parameter values listed in Table 1.

Applying our Bayesian framework to published datasets substantially reconciles estimates of cell-size regulation across organisms and growth conditions. In *E. coli* [14, 15, *16, 17, 24], S. pombe* [18], and several mammalian cell lines [7], conventional pooled estimates vary widely, whereas the Bayesian estimates are concentrated near the adder value (Fig. 4). The inferred noise amplitudes quantitatively account for the variation in pooled estimates across conditions through the opposing biases induced by extrinsic variability and measurement noise. We further apply the framework to *B. subtilis* [14, 25] *and Mycobacterium tuberculosis* [26], obtaining results broadly consistent with previously reported adder-like behavior. However, the mouse leukemia dataset [23] constitutes a notable exception, for which Bayesian inference yields 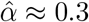, indicating a departure from adder-like regulation. Together, these results suggest that much of the apparent variation in cell-size regulation across organisms and growth conditions arises from estimator bias rather than changes in the underlying regulatory strategy, and most of the bacterial and eukaryotic datasets examined are compatible with *adder* -like regulation.

**Table 1:**
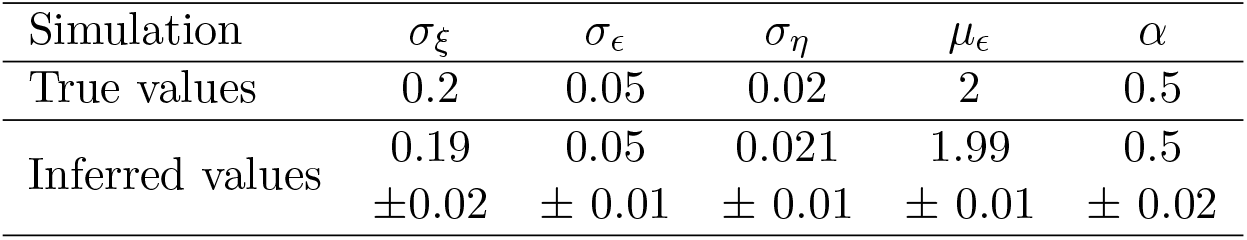
Validation of Bayesian inference on simulated data. True parameter values, *θ*, used to generate the simulated datasets, and the corresponding inferred values, 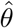, with 95% credible intervals obtained from the Bayesian analysis for *n*_*ℓ*_ = 15 and *N* = 50. The inferred parameters closely recover the true values, demonstrating the accuracy of the method on synthetic data.

**Figure 4:**
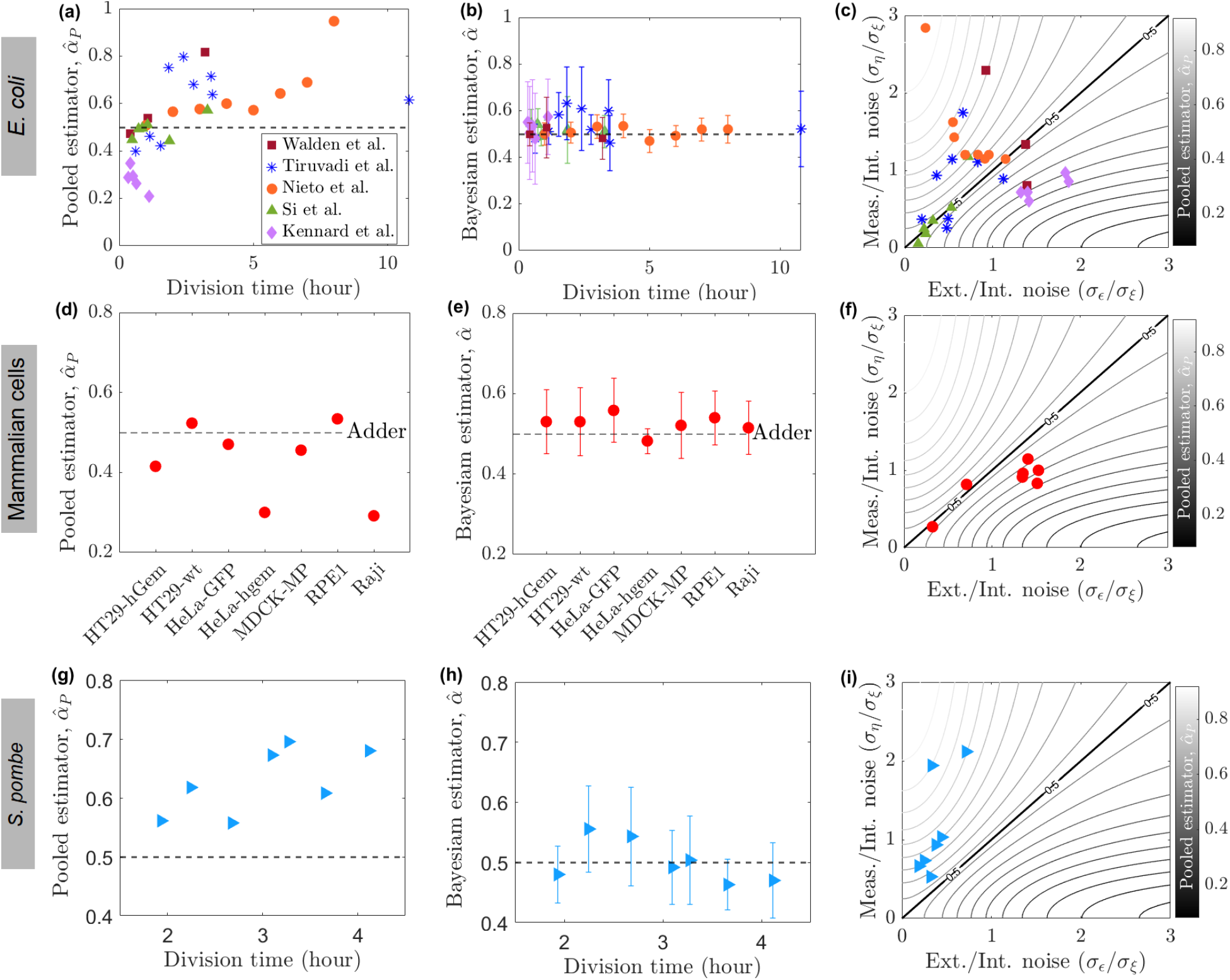
Bayesian inference shows adder cell-size regulation across organisms. Rows correspond to published datasets from (**a**–**c**) *E. coli* [14, 15, 16, 17, 24], (**d**–**f** ) mammalian cell lines [7], and (**g**–**i**) *S. pombe* across growth conditions [18]. (**a, d, g**) Conventional pooled estimates of the size-regulation parameter, 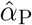, plotted against mean division time for *E. coli* and *S. pombe* and by cell line for mammalian cells. (**b, e, h**) Corresponding Bayesian estimates of *α*; symbols denote posterior means, and error bars denote 95% credible intervals. Horizontal dashed lines in the estimator panels mark the adder value, *α* = 0.5. (**c, f, i**) Each dataset is positioned according to its inferred extrinsic-to-intrinsic noise ratio, *σ*_*ϵ*_*/σ*_*ξ*_, and measurement-to-intrinsic noise ratio, *σ*_*η*_*/σ*_*ξ*_. The shading and contour lines show the analytical expectation of the pooled estimator from Eq. (9), assuming an underlying adder with *α* = 0.5. The thick black contour marks 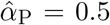, along which the opposing biases from extrinsic variability and measurement noise cancel. Above this contour, measurement noise dominates, biasing 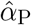 upward; below it, extrinsic variability dominates, biasing 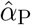 downward. Symbols and colors identify the corresponding datasets in the pooled-estimator panels. For each dataset, the value of 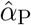 predicted by its location in the noiseratio plane closely agrees with the observed value in panels (**a, d, g**). Thus, differences in the relative amplitudes of measurement and extrinsic noise can account for the broad range of pooled estimates without requiring corresponding changes in the underlying regulation parameter.

## 3 Models

We consider a collection of cell lineages indexed by *ℓ*, with generations indexed by *t*. For a cell born with volume *v*_*ℓ,t*_, the division volume 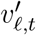 follows Eq. (1). The birth volume of the tracked daughter is related to the division volume of its mother by

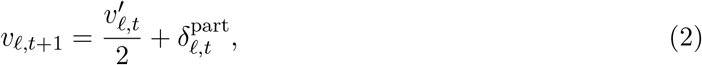

where 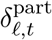 denotes the partitioning noise around symmetric division. Substituting Eq. (1) gives the following generation-to-generation dynamics for birth volume:

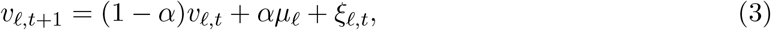

where *ξ*_*ℓ,t*_ is the effective intrinsic fluctuation that combines noise in division-size regulation with stochastic asymmetry in the partitioning of cellular material at division. We approximate the fluctuations *ξ*_*ℓ,t*_ as independent and identically distributed across lineages and generations, with zero mean and variance 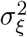:

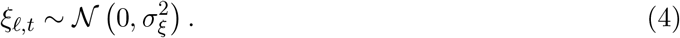

Fig. 1(a) shows the simulated birth and division sizes together with cell-size dynamics for a lineage associated with intrinsic noise only.

The birth-volume dynamics in Eq. (3) also extend to systematic asymmetric division, such as budding in yeast [4, 27]. If 1*/q* denotes the mean fraction of the division volume inherited by the daughter, then 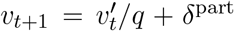. Symmetric division corresponds to *q* = 2, whereas *q >* 2 describes asymmetric division in which the daughter inherits less than half of the division volume. With this definition, daughter birth volumes in budding yeast follow the same generation-to-generation AR(1) dynamics as Eq. (3) (see Appendix Sec. D). However, the value of *α* that corresponds to an adder depends on *q*. The defining feature of an adder is the accumulation of constant volume from birth to division, leading to a unit slope when regressing the division volume against the birth volume. Here, this leads to the requirement *q*(1 − *α*) = 1, implying that *α* = 1 − 1*/q*. Thus, for adder, the familiar value *α* = 0.5 applies to symmetric division, whereas *α >* 0.5 for asymmetric division with *q >* 2. A detailed derivation for budding yeast is provided in Appendix Sec. D.

Measured cell sizes are subject to experimental noise. We model measurement noise as additive with mean 0 and variance 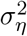. The measured birth size is therefore

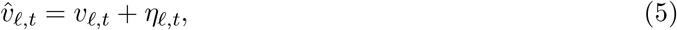

where

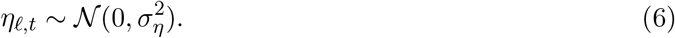

The measurement-noise terms are independent across lineages and generations. Fig. 1(b) shows the simulated measurements of birth and division sizes for a lineage in the presence of measurement noise that differs from the true ones.

In addition to fluctuations within individual lineages, single-cell datasets may contain persistent lineage-to-lineage differences in average cell size [28]. Such differences can arise from slowly varying physiological or environmental factors and cannot be represented by the generation-specific intrinsic noise. We therefore model extrinsic variability through the lineage-specific mean birth size,

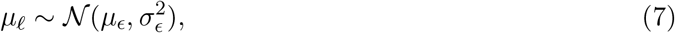

where *µ*_*ϵ*_ is the population-level mean birth size and 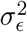 quantifies the variance in mean size across lineages. In this hierarchical formulation, *µ*_*ℓ*_ remains fixed within a lineage but varies across lineages. Extrinsic variability therefore shifts the characteristic size of an entire lineage, whereas intrinsic noise produces generation-to-generation fluctuations around that characteristic size. Fig. 1(c) illustrates two simulated lineages with different values of *µ*_*ℓ*_. For reference, the principal notation and model parameters used throughout are summarized in Appendix Table 2.

**Table 2:** Principal notation and model parameters. Quantities used in the cell-size model, Bayesian inference, and conventional estimators. Mean cell size has the units of the experimentally reported size variable; variances have squared size units.

| Symbol | Description |
| --- | --- |
| *Indices and data* |  |
| $\ell$ | Lineage index, $\ell = 1, \dots, N$. |
| $t$ | Generation index within a lineage, $t = 1, \dots, n_\ell$. |
| $N$ | Number of lineages in a dataset. |
| $n_\ell$ | Number of observed generations in lineage $\ell$. |
| $v_{\ell,t}$ | Latent (true) birth size of generation $t$ in lineage $\ell$. |
| $v'_{\ell,t}$ | Division size of generation $t$ in lineage $\ell$. |
| $\hat{v}_{\ell,t}$ | Measured birth size, $\hat{v}_{\ell,t} = v_{\ell,t} + \eta_{\ell,t}$. |
| *Model parameters and stochastic terms* |  |
| $\alpha$ | Dimensionless cell-size regulation parameter, assumed to be common to all lineages. The limits $\alpha \rightarrow 0^+$, $\alpha = 0.5$, and $\alpha = 1$ correspond to timer-like, adder, and sizer regulation, respectively. Stable size homeostasis requires $0 < \alpha < 2$. |
| $\mu_\ell$ | Mean birth size of lineage $\ell$; constant within a lineage and allowed to vary among lineages. |
| $\mu_\epsilon$ | Population mean of the lineage-specific mean birth sizes $\mu_\ell$. |
| $\sigma_\epsilon$ | Standard deviation of $\mu_\ell$ across lineages; amplitude of extrinsic lineage-to-lineage variability. |
| $\xi'_{\ell,t}$ | Zero-mean fluctuation in the division-size regulation equation. |
| $\delta_{\ell,t}^{\text{part}}$ | Deviation from equal partitioning of mother-cell size between the two daughter cells. |
| $\xi_{\ell,t}$ | Effective intrinsic fluctuation in the birth-size dynamics, combining division-regulation and partitioning noise; $\xi_{\ell,t} \sim \mathcal{N}(0, \sigma_\xi^2)$. |
| $\sigma_\xi$ | Standard deviation of the effective intrinsic fluctuations within lineages. |
| $\eta_{\ell,t}$ | Additive measurement error, $\eta_{\ell,t} \sim \mathcal{N}(0, \sigma_\eta^2)$. |
| $\sigma_\eta$ | Standard deviation of measurement noise. |
| $\sigma_{\text{int}}^2$ | Conditional stationary variance of latent birth size within a lineage, $\sigma_{\text{int}}^2 = \sigma_\xi^2 / [\alpha(2 - \alpha)]$. |
| $\theta$ | Full inferred parameter vector comprising $\mu_{1:N}$, $\alpha$, $\mu_\epsilon$, $\sigma_\epsilon$, $\sigma_\xi$, and $\sigma_\eta$, where $\mu_{1:N} = (\mu_1, \dots, \mu_N)$. |
| *Conventional estimators* |  |
| $\hat{\alpha}_P$ | Pooled estimate obtained from all lineage observations. |
| $\hat{\alpha}_\ell$ | Size-regulation estimate obtained separately within lineage $\ell$. |
| $\hat{\alpha}_{\text{LW}}$ | Lineage-wise estimate, mean of $\hat{\alpha}_\ell$. |

## 4 Results

### 4.1 Limitations of the existing methods

Two approaches in the literature are used to estimate the size-regulation parameter *α*. In the following section, we show that each estimator is affected by distinct systematic errors in the presence of measurement noise, extrinsic noise, and finite lineage length that are evident in single-cell data.

#### 4.1.1 Pooled estimator

A common approach to estimating *α* is to pool cell-size measurements across all lineages and infer a single effective size-regulation parameter from the combined dataset:

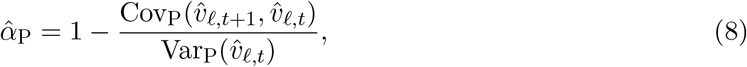

where Cov_P_ and Var_P_ denote the sample covariance and variance, respectively, evaluated over the pooled observations.

In the absence of measurement and extrinsic noise, the pooled estimator recovers the true value of *α* [Fig. 1(d)]. Independent measurement noise in birth size increases the variance of the observed birth sizes but leaves their covariance across successive generations unchanged. It therefore attenuates the pooled regression slope and biases 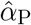 upward [Fig. 1(e)]. This attenuation of the regression slope toward zero is commonly referred to as regression dilution bias [30, 31]. By contrast, slowly varying extrinsic noise generates lineage-specific mean birth sizes, such that lineages with larger mean birth sizes also tend to have larger birth sizes in successive generations. Pooling across lineages therefore introduces a positive between-lineage covariance, increases the pooled regression slope, and biases 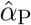 downward [Fig. 1(f)]. This discrepancy between the within-lineage regression slopes and the pooled regression slope is a form of aggregation bias related to Simpson’s paradox [29] [see Fig. 1(f)].

Indeed, the expected value of the pooled estimator can be obtained in closed form under the extrinsic-noise model in Eq. (7), the measurement-noise model in Eq. (5), and the size-regulation model in Eq. (1):

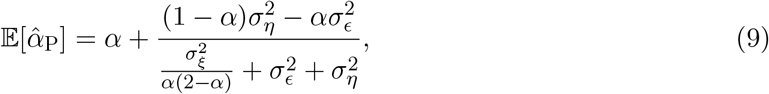

as detailed in Appendix Sec. A. The analytical prediction agrees with simulations for varying levels of measurement and extrinsic noise [Fig. 1(g)]. Fig. 1(h) shows a heat map of 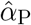 for the true value *α* = 0.5 in the plane of measurement and extrinsic noise, both normalized by the intrinsic noise. Measurement noise biases the pooled estimate upward, whereas extrinsic noise biases it downward. In the absence of both noise sources, the pooled estimator recovers the true value, as follows from Eq. (9). When both are present, the upward bias from measurement noise and the downward bias from extrinsic noise cancel when 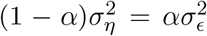, yielding pooled estimates equal to the true value. For *α* = 0.5, this condition reduces to *σ*_*η*_ = *σ*_*ϵ*_.

However, Eq. (9) cannot be used directly to recover the true *α* because the intrinsic (*σ*_*ξ*_), extrinsic (*σ*_*ϵ*_), and measurement (*σ*_*η*_) noise amplitudes are unknown from single-cell measurements.

#### 4.1.2 Lineage-wise estimator

To avoid the bias in the pooled estimator caused by lineage-to-lineage variation in mean cell size, a recent study [23] proposed estimating *α* separately for each lineage and then averaging the resulting estimates. We refer to this approach as the *lineage-wise estimator* :

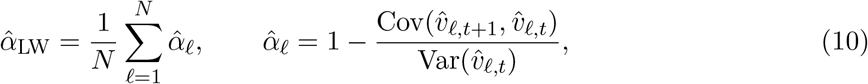

where *N* is the number of lineages, Cov and Var denote the sample covariance and variance, respectively, and 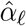 is the size-regulation parameter estimated within lineage *ℓ*.

For short lineages, the estimated regression slopes between birth sizes in successive generations (equivalently, between birth and division size) can vary substantially across lineages. Figs. 2(a) and 2(b) show representative cell-size trajectories and the corresponding lineage-specific regression estimates, respectively. The variance of 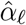 is expected to scale approximately as 1*/*(*n*_*ℓ*_−1) (*n*_*ℓ*_ is the length of lineage *ℓ*) according to standard asymptotic approximation or the central limit theorem, as 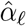 are independent across lineages (Fig. 2(c)). The analytical expression for the variance of 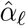 is derived in Appendix Sec. B in Eq. (B.55). Less intuitively, finite lineage length also biases the mean of 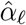, as shown in Fig. 2(d). This bias arises because 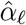 depends on the ratio of the sample covariance to the sample variance. The generation-to-generation birth-size recursion in Eq. (3) is equivalent to a first-order autoregressive mode, with autoregressive coefficient 1 − *α*. The resulting bias therefore corresponds to the classical finite-sample bias of autoregressive coefficient estimators [33, 34].

Independent measurement noise introduces an additional upward bias in the lineage-wise estimator through regression dilution, and this bias persists as *n*_*ℓ*_ → ∞. Combining the measurement-noise bias with the leading-order finite lineage length correction gives

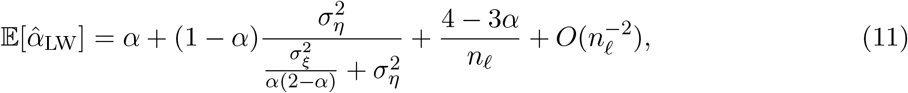

as derived in Appendix Sec. B. Simulations confirm the decay of the finite-sample bias with lineage length and the persistent upward shift caused by measurement noise [Fig. 2(e)]. A similar dependence on lineage length is observed in single-cell data from *E. coli* [32] and mouse leukemia cells [23] [Figs. 2(f) and 2(g), respectively]. For the *E. coli* dataset, each of the 54 lineages, originally spanning 70 generations, was truncated to a range of lengths, and the lineage-wise estimates were averaged across lineages at each truncation length. For the mouse leukemia dataset, the 24 lineages span 3–29 generations; the individual estimates 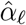 were therefore grouped by lineage length and plotted together with the corresponding binned averages.

However, although the lineage length *n*_*ℓ*_ is known from the data, Eq. (11) cannot be used directly to recover the true *α* because the measurement-noise amplitude *σ*_*η*_ is unknown.

## 5 Methods: Bayesian inference

The pooled and lineage-wise estimators introduced above can both yield biased estimates of *α*. Although the relationships between these estimates and the true regulation parameter can be derived analytically (in Eqs. (9) and (11)), they depend on the unknown amplitudes of intrinsic, extrinsic, and measurement noise and therefore cannot be inverted directly to recover the true value of *α*. We therefore adopt a Bayesian framework to jointly infer *α*, the lineage-specific mean birth volumes, and the noise parameters from the posterior distribution conditioned on the observed lineage data.

For the model introduced in Sec. 3, we perform Bayesian inference for the unknown parameters

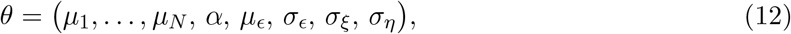

given the observed lineage data

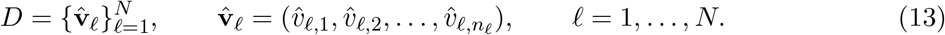

The lineage-specific parameters *µ*_*ℓ*_ account for variation in average birth size across lineages. Such variation may arise from extrinsic factors, including fluctuations in growth rate. This interpretation is consistent with the Schaechter–Maaløe–Kjeldgaard growth law, which describes an approximately exponential dependence of average cell size on growth rate under balanced-growth conditions [35]. By contrast, the size-regulation parameter *α* is taken to be common to all lineages, consistent with the assumption that the underlying size-regulation strategy remains unchanged across lineages. The posterior distribution is

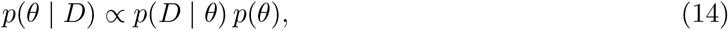

where *p*(*D* | *θ*) denotes the observed-data likelihood and *p*(*θ*) the prior distribution. Although the observed process 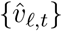 is non-Markovian, the model admits a linear-Gaussian state-space representation. Consequently, the exact likelihood can be evaluated recursively using the Kalman filter via the one-step-ahead predictive densities 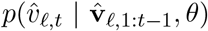 [36, 37]. Posterior inference is performed using Markov chain Monte Carlo (MCMC). Details of the derivation are provided in Appendix C. We report posterior means as point estimates with 95% credible intervals, defined by the 2.5th and 97.5th percentiles of the corresponding marginal posterior distributions.

### 5.1 Synthetic data

To validate the Bayesian estimator, we apply it to simulated cell size data obtained from the model in Sec. 3 and compare the inferred parameter values with the true ones. Results for 50 lineages of length 15 generations are summarized in Table 1. In all cases, the true parameter values lie within the corresponding 95% credible intervals. Moreover, Figs. 3(a)–3(c) show the marginal posterior distributions of the size-regulation parameter for timer-like (*α* = 0.1), adder (*α* = 0.5), and sizer (*α* = 1) regimes, respectively. In all three cases, the Bayesian estimate accurately recovers the true value of *α*. Fig. 3(d) shows that the Bayesian estimator remains centered on the true value in the presence of measurement and extrinsic noise, whereas the pooled estimator 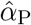 is biased.

Fig. 3(e) shows that the Bayesian estimator also recovers the true value for short lineages, whereas the lineage-wise estimator 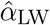 is biased upward at small lineage lengths.

To assess the strength of evidence supporting the inferred size-regulation parameter *α*, we complement Bayesian inference with the Bayes factor (BF) [38] (see Appendix Sec. C.6). The BF quantifies the relative evidence for competing models through the ratio of their marginal likelihoods, with a BF of 10 indicating strong evidence for the pure adder model (*α* = 0.5) over mixed strategies (*α* ≠ 0.5). As shown in Fig. 3(d) and (e) insets, where synthetic data were generated with *α* = 0.5, the Bayesian estimates accurately recover the true parameter value, while the BFs provide complementary evidence favoring the pure adder model, even for short lineages and in the presence of extrinsic and measurement noise.

Having established the reliability of the Bayesian estimator, we next apply it to experimental datasets and compare the inferred values of *α* with those obtained using previously reported pooled estimators.

### 5.2 Experimental data

We first revisit published lineage-resolved measurements of *E. coli* cell size spanning a broad range of growth conditions. Most of these datasets were acquired using microfluidic mother-machine devices, which enable high-throughput tracking of individual cells over many generations under controlled growth conditions [3, 16, 14, 17, 24, 12]. The dataset reported by Kennard *et al*. [15] was instead obtained by time-lapse tracking of growing microcolonies on agarose pads. After processing, each condition contains up to approximately 10^3^ short lineages, typically spanning three to four generations.

The conventional pooled estimates vary substantially across datasets, spanning timer-like to sizer-like regulation across growth conditions, consistent with previous reports [Fig. 4(a)]. By contrast, the Bayesian estimates cluster near the adder value, *α* = 0.5, across the full range of growth rates, from fast to slow, with uncertainty quantified by the 95% posterior credible intervals [Fig. 4(b)]. The deviations of the pooled estimates from the adder value can be understood by differences in the inferred noise amplitudes. When the datasets are positioned in the plane of extrinsic-to-intrinsic noise, *σ*_*ϵ*_*/σ*_*ξ*_, and measurement-to-intrinsic noise, *σ*_*η*_*/σ*_*ξ*_, their pooled estimates closely follow the analytical contours predicted by Eq. (9) for an underlying adder with true *α* = 0.5 [Fig. 4(c)]. The diagonal *σ*_*ϵ*_ = *σ*_*η*_ separates the two bias regimes: measurement-noise-dominated datasets yield upward-biased pooled estimates, whereas extrinsic-noise-dominated datasets yield downward-biased estimates.[Fig. 4(c)].

We next apply the same analysis to published lineage-resolved measurements from mammalian cell lines [7]. In this study, cell volume was tracked over complete division cycles using fluorescence-exclusion microscopy and, for selected cell lines, microchannels of known cross-sectional area, in which volume was inferred from cell length. The conventional pooled estimates vary widely across cell lines and deviate substantially from the adder value [Fig. 4(d)]. By contrast, the Bayesian estimates cluster near the adder value, *α* = 0.5, with uncertainty quantified by the 95% posterior credible intervals [Fig. 4(e)]. As in *E. coli*, the deviations of the pooled estimates from the adder value are explained by the inferred ratio between extrinsic and measurement noise [Fig. 4(f)].

For fission yeast, we analyzed the long-term single-cell trajectories of *S. pombe* [18] in different culture media and temperatures. In these experiments, individual old-pole cells were retained in a microfluidic mother-machine device under continuous medium perfusion for up to 50–80 generations, with cell size quantified from projected cell area. The conventional pooled estimates lie systematically above the adder value, *α* = 0.5, ranging from approximately 0.56 to 0.70 and thus suggesting regulation intermediate between an adder and a sizer [Fig. 4(g)] consistent with the previous studies [20]. By contrast, the Bayesian estimates cluster near *α* = 0.5, with the adder value contained within the 95% posterior credible intervals for all seven conditions [Fig. 4(h)]. The apparent sizer-like shift of the pooled estimates is explained by the inferred noise amplitudes: all datasets lie in the measurement-noise-dominated regime, *σ*_*η*_ *> σ*_*ϵ*_, and closely follow the analytical contours predicted by Eq. (9) for an underlying adder [Fig. 4(i)]. Thus, after accounting explicitly for intrinsic, extrinsic, and measurement noise, fission-yeast size regulation is adder-like across different growth conditions.

We further apply the Bayesian framework to published datasets for *B. subtilis* [14] for several growth conditions (see Appendix Fig. C.1), *M. tuberculosis* [26], and mouse leukemia cells [23]. The inferred parameters are listed in Appendix Table 3. For *B. subtilis*, and *M. tuberculosis*, the inferred regulation is adder-like, consistent with previous results obtained using pooled estimators. By contrast, the mouse leukemia cells show a departure from adder-like regulation with inferred 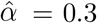.

**Table 3:**
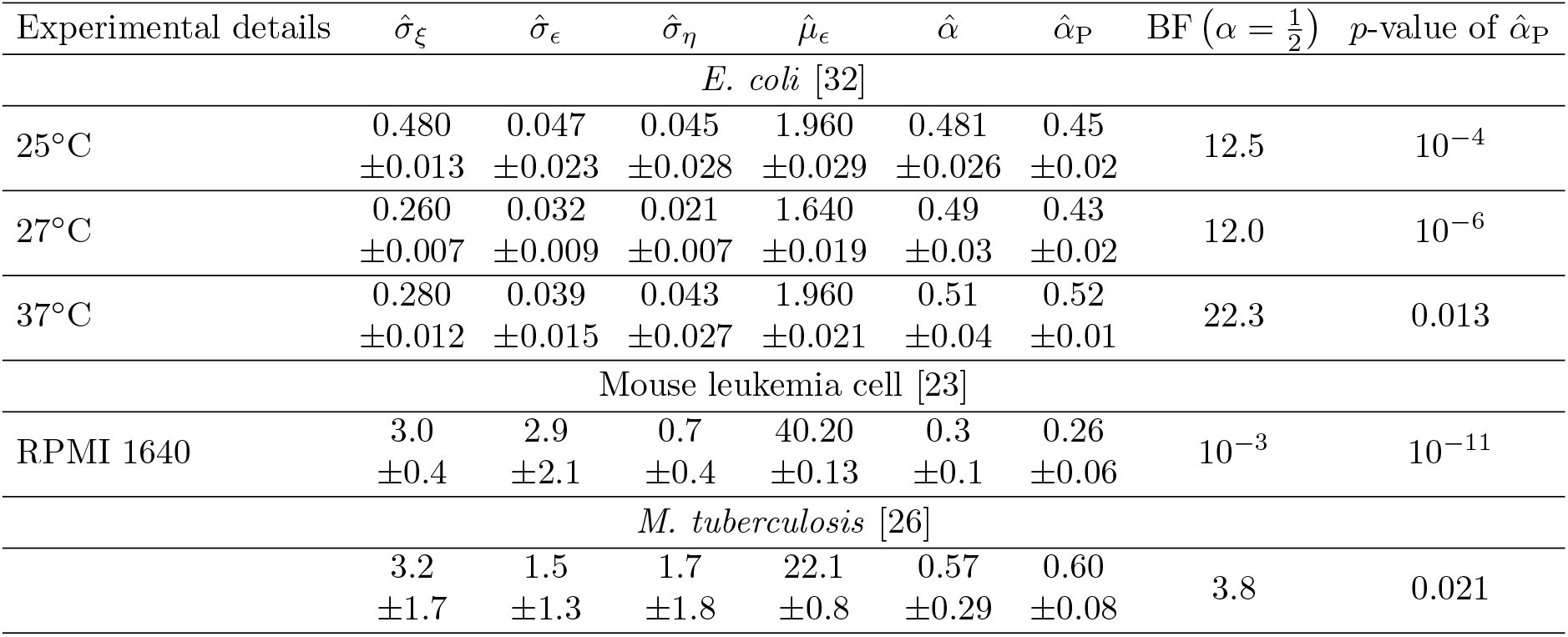
Inferred parameters 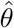 for the experimental datasets. Here, *σ*_*ξ*_, *σ*_*ϵ*_, and *σ*_*η*_ denote the standard deviations of intrinsic, extrinsic (between-lineage), and measurement noise, respectively; *µ*_*ϵ*_ is the population mean of the lineage-specific mean birth sizes *µ*_*ℓ*_; and *α* is the size-regulation parameter. The Bayes factor is defined as BF_12_ = *p*(*D* | *M*_1_)*/p*(*D* | *M*_2_), where *M*_1_ is the adder model with *α* = 0.5 and *M*_2_ is the general model with *α* ∼ *U*(0, 2). Following Jeffreys’ scale [48], BF_12_ ≥ 3.2 is interpreted as substantial evidence for the adder model. The reported *p*-value is obtained from a two-sided *t*-test of the null hypothesis *H*_0_ : *α* = 0.5 against *H*_1_ : *α* ≠ 0.5, based on the conventional pooled estimate 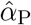 and its standard error. A *p*-value below 0.05 indicates a statistically significant deviation from the adder value, whereas a larger *p*-value does not, by itself, establish adder regulation. Bayesian estimates are reported with 95% credible intervals, whereas the conventional pooled estimates are reported with 95% confidence intervals.

Together, these results indicate that the apparent variation in size-regulation behavior across bacterial and mammalian datasets can largely arise from noise-induced bias in the pooled estimator. After accounting for intrinsic, extrinsic, and measurement noise, the Bayesian analysis supports adder size regulation across most datasets considered from bacteria to eukaryotes except one mouse leukemia cell dataset.

## 6 Discussion

In this study, we showed that two widely used estimators of the cell-size regulation parameter *α* can be substantially biased. The lineage-wise estimator 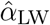 is biased upward for short lineages, whereas the pooled estimator 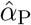 is biased downward by lineage-to-lineage variability in mean cell size. Measurement noise introduces an additional upward bias in both estimators. These predictions are consistent with experimental data of bacteria and eukaryotic cells. To address these limitations, we developed a Bayesian framework that jointly infers the size-regulation parameter, the amplitudes of intrinsic, extrinsic, and measurement noise, and the mean birth size. Reanalysis of the experimental data supports adder-like regulation in most datasets considered, including *E. coli* across a broad range of growth conditions and *S. pombe* across different culture media and temperatures, while retaining a deviation from adder-like regulation in mouse leukemia cells.

The estimator biases identified here have direct implications for comparisons of cell-size regulation across experiments. Variation in conventional estimates of *α*, including apparent transitions between timer-, adder-, and sizer-like behavior, does not necessarily reflect a change in the underlying biological mechanism. Instead, part of this variation can arise from differences in lineage length, measurement precision, and lineage-to-lineage heterogeneity. Comparisons of *α* across organisms or experimental conditions should therefore account explicitly for these features of the data. Within the datasets analyzed here, the Bayesian framework explains much of the variation in pooled estimates through differences in the relative amplitudes of extrinsic and measurement noise. This result indicates that most previously reported deviations from adder-like regulation may result from statistical bias rather than biological variation.

Our results are based on the stochastic model in Eq. (1), which describes division size as a linear function of birth size with additive noise. The linear relation represents a first-order approximation of a general size-regulation function around the mean birth size. Higher-order terms describe nonlinear behavior away from the mean but do not alter the definition of *α* as the first-order regulation parameter [8, 10]. For narrow size distributions, these nonlinear contributions are expected to be small, and previous theoretical and empirical studies have shown that the linear approximation accurately describes bacterial cell-size data [8, 10, 11]. Models that are linear in logarithmic size also reduce to the same first-order description when size fluctuations are small [8, 9, 10, 11]. For broader size distributions, higher-order terms may be required to describe the full dependence of division size on birth size, while *α* retains its interpretation as the local first-order regulation parameter.

Our model assumes one-generation mother–daughter dependence along individual lineages. Recent work using Bayesian inference on a stochastic division-rate model found that one-generation memory is sufficient to describe the bacterial and eukaryotic data [39]. Nevertheless, other systems exhibit dynamics beyond simple one-generation dependence, including strong cousin–cousin correlations in mammalian lineage trees and circadian modulation of cell-cycle timing in cyanobacteria [40, 41]. The present framework does not include longer-range intergenerational memory, correlations within branching pedigrees such as sister–sister and cousin–cousin correlations, temporal correlations in the noise parameters, or time-dependent modulation by endogenous oscillators. Extensions to nonlinear or higher-order temporal dynamics and complete lineage trees would allow the robustness of the inferred regulation parameter to be tested beyond the assumptions considered here.

More broadly, the Bayesian framework can be extended beyond the inference of a single size-regulation parameter to investigate how cell size is coordinated with growth rate, division timing, and other cellular variables. For example, Cadart *et al*. [7] examined the relative contributions of growth rate and cell-cycle duration using regression-based analyses. Because such estimates may also be affected by measurement noise, finite sampling, and cell-to-cell heterogeneity, the corresponding models could be reformulated within the Bayesian framework to jointly infer regulatory couplings and noise amplitudes. Such extensions would allow more robust tests of how growth and division jointly determine cell size and provide a natural direction for future work.

More fundamentally, the prevalence of adder-like regulation across the diverse datasets examined here raises an evolutionary question: why does the adder recur across phylogenetically distant organisms? One proposed explanation is that adder control minimizes size fluctuations generated by intrinsic molecular noise [21]. Recent work, however, showed that this minimum-noise property does not generally persist when extrinsic threshold fluctuations and additional mechanistic factors are incorporated [42]. Other possible explanations include fitness advantages under particular selective regimes [43] and constraints imposed by cell-cycle architecture [44]. Determining whether this recurrence reflects selection for enhanced fitness or robustness to molecular and environmental perturbations, or instead emerges from general features of cell-cycle architecture, therefore remains a central challenge for future work.

## Code availability

The code implementing the Bayesian framework presented in this study is publicly available on GitHub in a toolbox named *Bayesian Inference of Size Control Using Independent Time-series* (BISCUIT) [45].

## Acknowledgments

This work was supported by the European Union (ERC, BIGR, 101125981) and the Israeli Science Foundation (146873). AA thanks the generous support of the Helen and Martin Kimmel Award for Innovative Investigation and the Clore Center for Biological Physics.

## A Estimation of bias in pooled estimator

In this SI Appendix, we derive Eq. (9) in the main text for the pooled estimator in the presence of lineage-to-lineage variability in mean birth size and measurement noise. The generation-to-generation dynamics of birth sizes in Eq. (3) of the main text follow a first-order autoregressive process, AR(1):

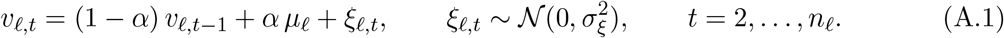

The within-lineage variance of birth-size fluctuations from Eq. (A.1) is given as

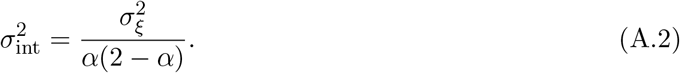

To represent slowly varying physiological or environmental factors, we assume that their effects are approximately constant within each lineage but differ among lineages. We therefore allow the lineage-specific mean birth size *µ*_*ℓ*_ to vary across lineages and quantify this extrinsic variability by

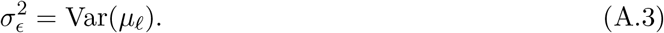

The measured birth size is

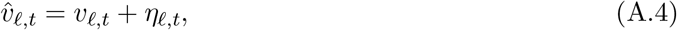

where *η*_*ℓ,t*_ is zero-mean measurement error with variance 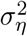. We assume that the measurement errors are independent and identical for each generation.

We first evaluate the pooled variance of the measured birth size by considering *v*_*ℓ,t*_ = *µ*_*ℓ*_ + *δ*_*ℓ,t*_:

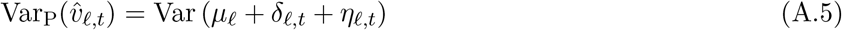

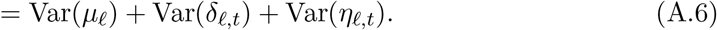

The cross terms vanish because E[*δ*_*ℓ,t*_ | *ℓ*] = 0 and the measurement noise is independent of the underlying cell sizes. Therefore,

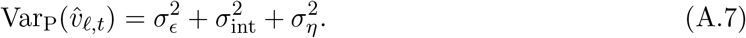

The pooled covariance between measured birth sizes in successive generations is:

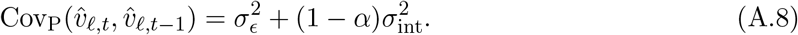

Thus, independent measurement noise in cell size does not affect the population covariance, although it increases the variance. Using Eqs. (A.7) and (A.8) the pooled estimator is given as follows:

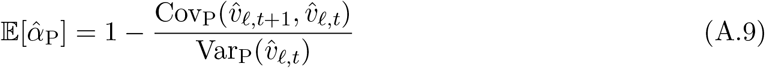

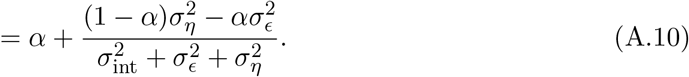

Finally, substituting Eq. (A.2) gives

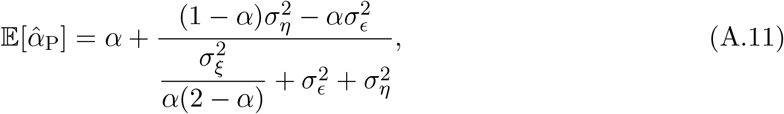

which is the large-sample prediction for the pooled estimator in the presence of both extrinsic and measurement noise.

Equation (9) in the main text shows that the two noise sources bias the pooled estimator in opposite directions. Lineage-to-lineage variability in mean birth size biases the estimate downward, whereas measurement noise biases it upward. In the absence of both noise sources, the true value *α* is recovered. When extrinsic noise dominates, the pooled estimate approaches zero; when measurement noise dominates, it approaches one. The two biases cancel when

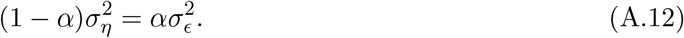

## B Effect of short lineage on the lineage-wise estimator

### B.1 Finite-sample bias of the lineage-wise AR(1) estimator

The generation-to-generation dynamics of birth sizes follow a first-order autoregressive process, AR(1), given in Eq. (3) in the main text. Defining the centered variable

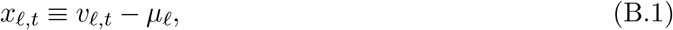

the dynamics become

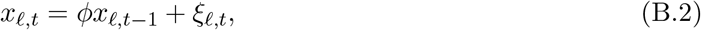

where

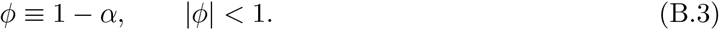

Thus, estimating *α* is equivalent to estimating the AR(1) slope *ϕ*.

We assume that each lineage is sampled from the stationary distribution of the AR(1) process. Then the moments of the latent variable *x*_*ℓ,t*_ will be as follows

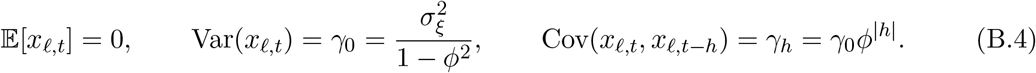

The usual estimated slope of the AR(1) for each lineage is

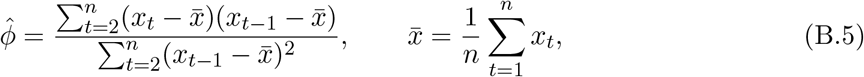

and the lineage-wise estimator will be

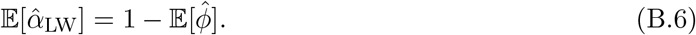

Please note that for notational simplicity, in the derivation below we suppress the lineage index *ℓ* and write *n* = *n*_*ℓ*_ and 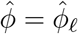.

#### B.1.1 Step 1: Bias in lineage-wise estimator when the mean is known

Here we will analyze the lineage-wise estimator with known mean. For this case, the estimated slope for each lineage in Eq. (B.5) will reduce to the following form

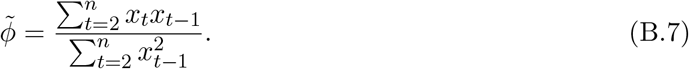

Using the relation, *x*_*t*_ = *ϕx*_*t−*1_ + *ξ*_*t*_, the numerator is given by

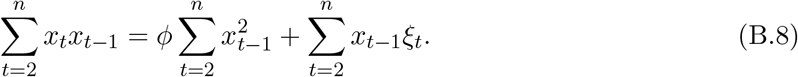

Therefore

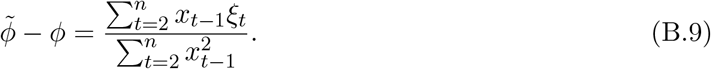

Taking the expectation of the preceding equation and expanding the denominator around the mean, 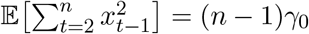 gives

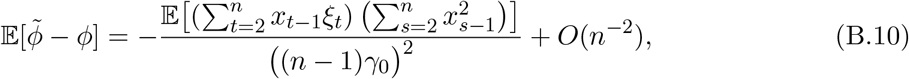

where we use the fact that E[*x*_*t−*1_*ξ*_*t*_] = 0. Now we need to solve

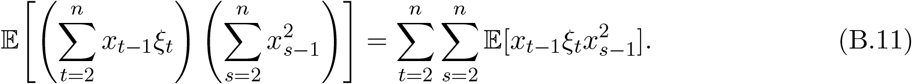

If *s* ≤ *t*, then 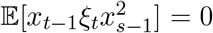. Hence only *s > t* contributes. Writing *h* = *s* − *t* ≥ 1, the AR(1) recursion gives

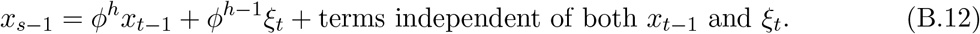

Squaring this expression and multiplying by *x*_*t−*1_*ξ*_*t*_, the only non-vanishing contribution after taking expectation is

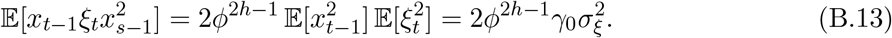

Therefore

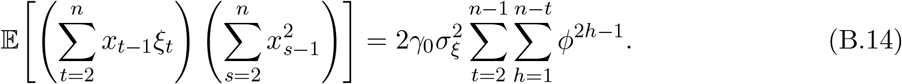

Using

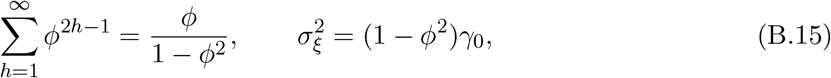

we obtain, to leading order in *n*,

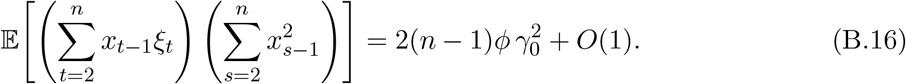

Substituting back,

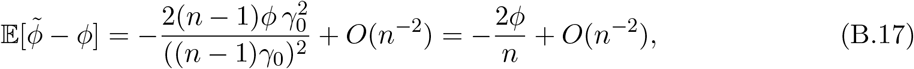

and hence

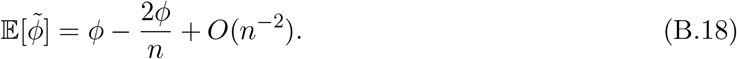

#### B.1.2 Step 2: Correction in lineage-wise estimator due to estimating the mean

When the mean is estimated from the data, the slope estimator for an individual lineage, 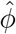, is given by Eq. (B.5). Expanding the numerator and denominator to first order in 1*/n* gives

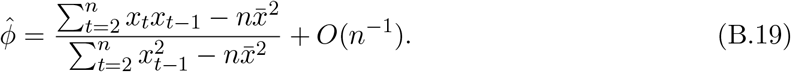

Expanding the above ratio around the known-mean estimator 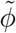 defined in Eq. (B.7) yields

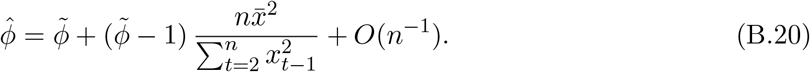

Taking expectations, and replacing 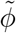 by *ϕ* inside the correction term (which changes the result only at order *O*(*n*^*−*2^)), we obtain

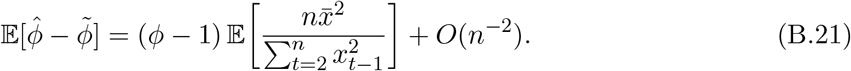

Now 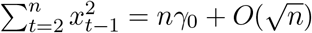, so

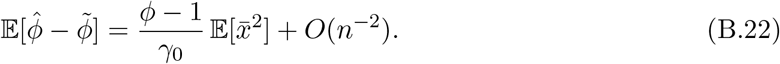

It remains to compute 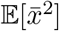. Since 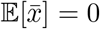,

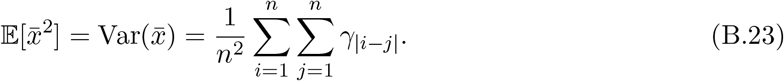

Grouping terms by lag *h* = |*i* − *j*|,

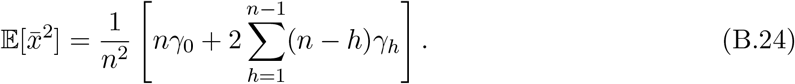

Using *γ*_*h*_ = *γ*_0_*ϕ*^*h*^ and simplifying, we obtain

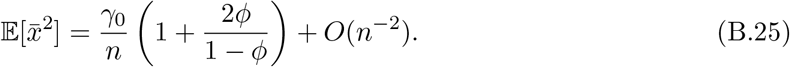

Combining Eq. (B.22) with the known-mean result in Eq. (B.18) we obtain

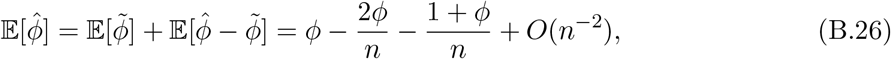

hence

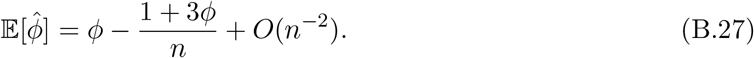

Finally, since 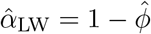 and *ϕ* = 1 − *α*,

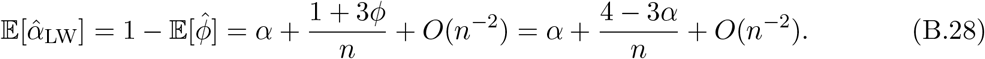

Restoring the lineage index *n* = *n*_*ℓ*_, we arrive at

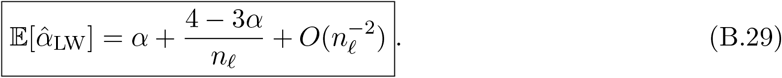

#### B.1.3 Explicit reduction of the Kendall–Marriott–Pope formula to the lineage-wise estimator

For completeness, we make explicit the connection between the lineage-wise estimator and the classical Kendall–Marriott–Pope result for the sample autocorrelation of an AR(1) process [33, 34]. For an AR(1) process with an estimated mean, the classical result for the lag-*k* sample autocorrelation *r*_*k*_, based on a sample of size *n*, is

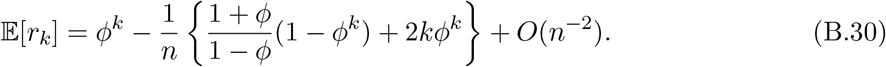

Setting *k* = 1 gives

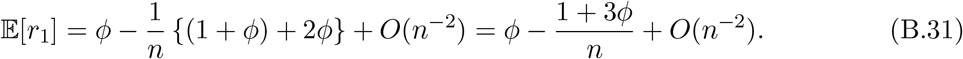

Identifying the lineage-wise estimator with the lag-one sample autocorrelation, 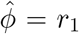, therefore recovers (B.27), and hence (B.29).

### B.2 Additional bias from measurement noise

In this section, we derive the effect of measurement noise on *α* analytically. For the latent dynamics of cell size in Eq. (3) in the main text, we model the observed variables as

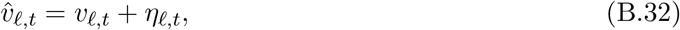

where the measurement errors *η*_*ℓ,t*_ are independent and identically distributed Gaussian random variables with mean zero and variance 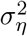, and are assumed to be independent of *v*_*ℓ,t*_ and *ξ*_*ℓ,t*_. For this model, the estimated *α* will be

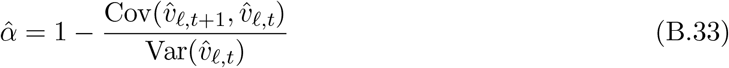

We now compute the covariance and variance entering the estimator. First,

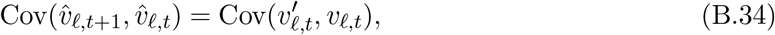

because all cross-terms involving the measurement errors vanish due to independence and zero mean. Therefore,

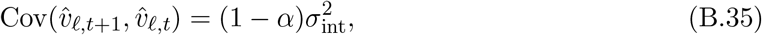

where

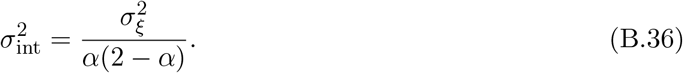

In contrast, the variance of the observed predictor is inflated by the measurement noise:

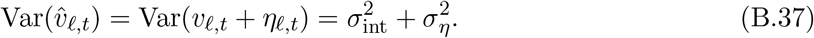

Now, substituting Eqs. (B.35) and (B.37) into Eq. (B.33) we obtain

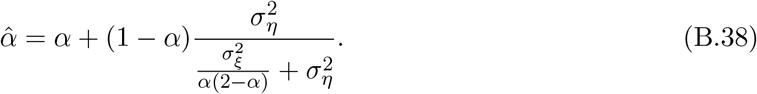

### B.3 Finite-sample variance of the lineage-specific estimator

We next derive the leading-order sampling variance of the lineage-specific estimator 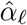 in the absence of measurement noise. As before, we suppress the lineage index *ℓ* and write 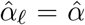 and *n*_*ℓ*_ = *n*. Similar to before, defining

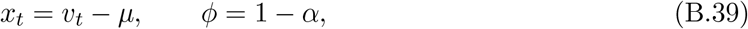

the birth-volume becomes AR(1) process as below

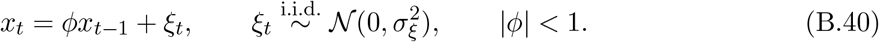

We assume that the process is stationary, so that

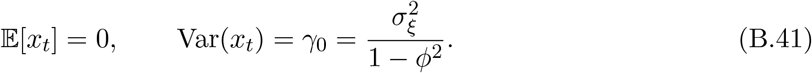

The lineage-specific slope estimator of the AR(1) process in Eq. (B.40) when the mean is estimated is given in Eq. (B.20). Substituting Eq. (B.9) into this expression yields

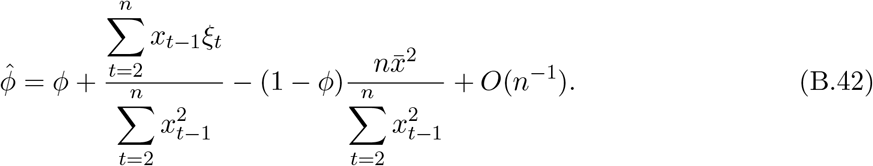

For a stationary and ergodic process, the sample average of 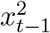 converges to its stationary expectation as the lineage length increases. Therefore, for sufficiently long lineages,

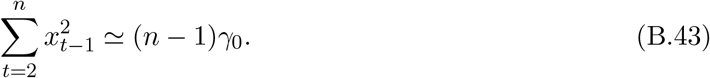

Moreover, combining 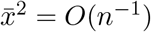 with Eq. (B.43) the following term in Eq. (B.42) becomes

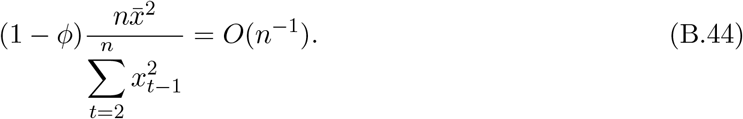

Consequently, Eq. (B.42) becomes

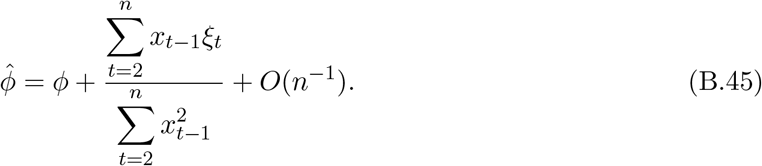

We next consider the numerator in the preceding equation

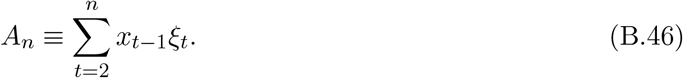

Since *x*_*t−*1_ and *ξ*_*t*_ are independent and E[*ξ*_*t*_] = 0, we have E[*A*_*n*_] = 0. The variance of *A*_*n*_ is

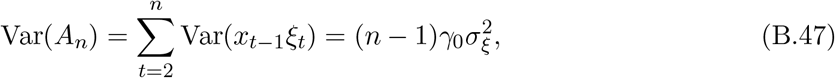

as all cross-covariance terms in the variance of *A*_*n*_ vanish. For any two generations *s* < *t*, the covariance between the corresponding numerator terms is

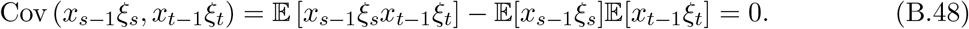

Since *x*_*s−*1_ and *ξ*_*s*_ are independent and have zero means, the second term,

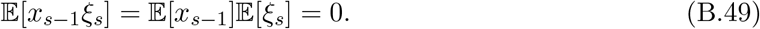

Moreover, for *s* < *t*, the quantities *x*_*s−*1_, *ξ*_*s*_, and *x*_*t−*1_ are all determined by the history of the process before *ξ*_*t*_. Since *ξ*_*t*_ is independent of this history and has zero mean, the first term,

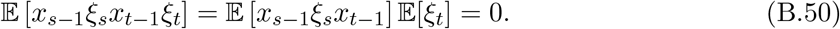

The central limit theorem for sufficiently large *n*, therefore gives

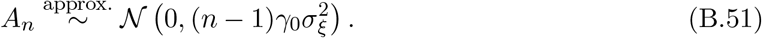

Combining (B.45) with Eqs. (B.43) and (B.51) gives

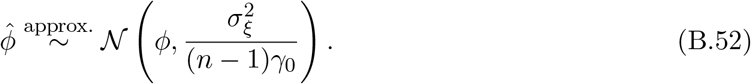

The leading-order variance of 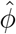 is therefore

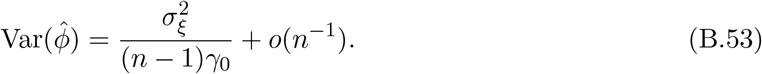

Using (B.41), and the relation *ϕ* = 1 − *α*, we obtain

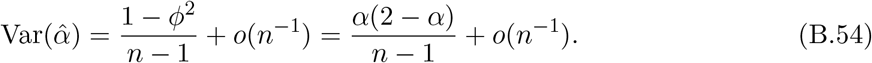

Restoring the lineage index gives

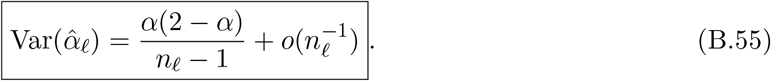

## C Bayesian inference

### C.1 Data structure and inferred parameters of the cell-size regulation model

We consider *N* independent cell lineages. For lineage *ℓ*, we observe a time series of measured birth sizes,

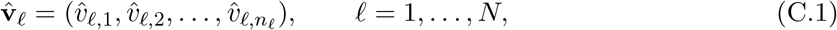

where *n*_*ℓ*_ denotes the number of generations in lineage *ℓ*. The lineage lengths may vary across lineages. The complete dataset is therefore

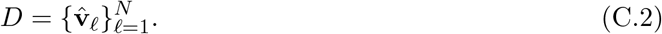

The measured birth size 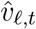 is assumed to be related to the latent birth size *v*_*ℓ,t*_ through additive Gaussian measurement noise,

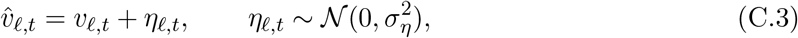

where 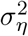 denotes the measurement-error variance. This term may incorporate multiple sources of observational uncertainty, including errors in size estimation and in the measurement of division time.

Within each lineage *ℓ*, the latent birth size *v*_*ℓ,t*_ is assumed to follow a first-order autoregressive process,

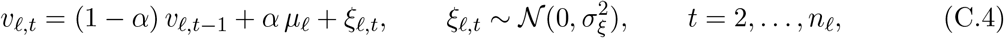

where *µ*_*ℓ*_ is the lineage-specific mean birth size, *α* is the size-regulation parameter, and 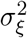 quantifies intrinsic biological variability within a lineage.

The unknown parameter vector for our model is

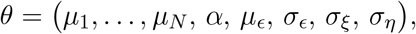

where (*µ*_*ϵ*_, *σ*_*ϵ*_) characterize the across-lineage distribution of the lineage-specific mean birth sizes, which are assumed to follow

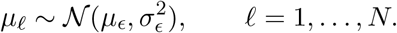

The variability of lineage-specific mean birth sizes arises from the extrinsic noise sources.

### C.2 Likelihood

In this section, we derive the exact likelihood for the observed data in the latent AR(1) model with additive Gaussian measurement noise. By the chain rule of probability, the exact observed-data likelihood is

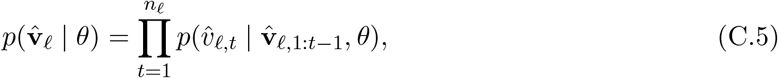

where 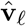 is defined in Eq. (C.1), and 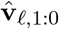 denotes the empty history. It is important to note that in contrast to the latent process, the observed process is not a first-order Markov process. Equivalently,

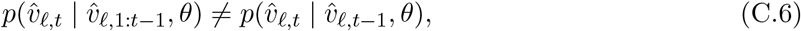

but, the latent process {*v*_*ℓ,t*_} in Eq. (C.4) is first-order Markov:

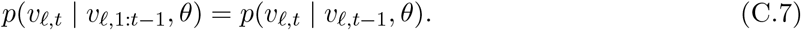

One can show the non-Markovianity of the observed variable by using 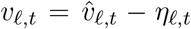, and substitution into Eq. (C.4) as follows

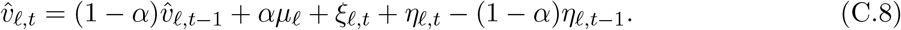

The effective noise term in the observed process is *ξ*_*ℓ,t*_ + *η*_*ℓ,t*_ − (1 − *α*)*η*_*ℓ,t−*1_, which is serially correlated if *α* ≠ 1 and 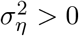.

Since the latent states are unobserved, each factor in the chain-rule in Eq. (C.5) is obtained by marginalizing over the latent state:

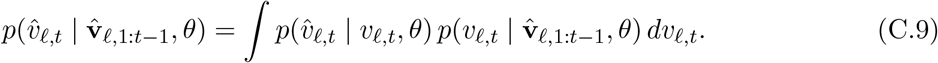

The first term is the observation density,

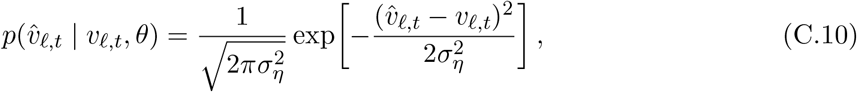

whereas the second term is the one-step predictive density of the latent state given the past observations,

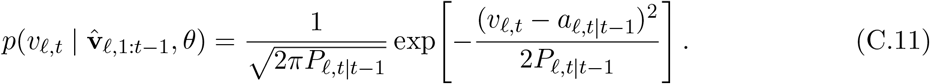

Here,

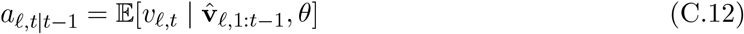

and

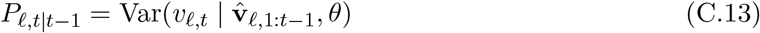

denote the one-step predictive mean and variance of the latent state, respectively. Eq. (C.11) remains Gaussian at every time step as both the latent and the observation equations are linear, and because both intrinsic noise *ξ* and measurement noise *η* are Gaussian. Consequently, the integral in Eq. (C.9) is a Gaussian convolution, and the predictive distribution of the observation is

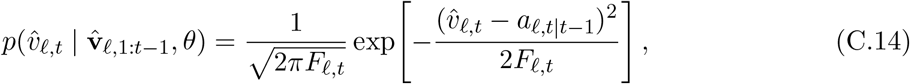

with

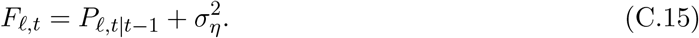

The quantities *a*_*ℓ,t*|*t−*1_, *P*_*ℓ,t*|*t−*1_, and hence *F*_*ℓ,t*_, are computed recursively by the Kalman filter [36, 37]. Under the stationary assumption, the initial predictive distribution of the latent state is

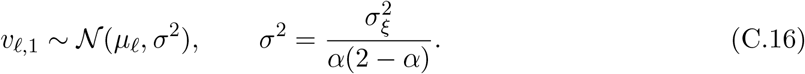

Equivalently, the Kalman filter is initialized with

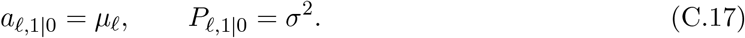

Starting from the stationary initialization in Eq. (C.16), the Kalman filter proceeds by alternating an update step and a prediction step. After observing 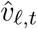, the filtered mean and variance of the latent state are updated as

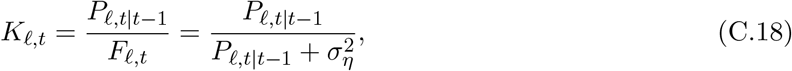

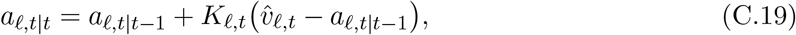

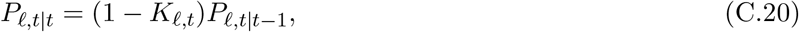

where *K*_*ℓ,t*_ is the Kalman gain. The mean update has the standard innovation form: the filtered mean equals the predictive mean plus a correction proportional to the prediction error 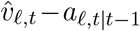. The proportionality factor *K*_*ℓ,t*_ determines the relative weight assigned to the new observation and the prior prediction. In particular, when the measurement noise is large relative to the predictive uncertainty, 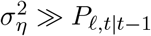, one has *K*_*ℓ,t*_ ≈ 0, so the observation contributes little to the update and *a*_*ℓ,t*|*t*_ ≈ *a*_*ℓ,t*|*t−*1_. Conversely, when the measurement noise is small, 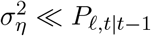, one has *K*_*ℓ,t*_ ≈ 1, so the filtered mean is pulled strongly toward the observation. The variance update reflects the same principle: conditioning on the new observation reduces uncertainty, so that *P*_*ℓ,t*|*t*_ ≤ *P*_*ℓ,t*|*t−*1_. The reduction is larger (smaller) when the measurement noise is smaller (larger).

The corresponding one-step prediction for the next latent state follows from the AR(1) dynamics in Eq. (C.4):

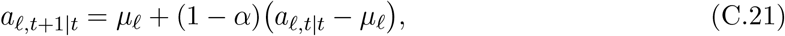

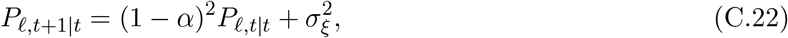

where we use the definition in Eqs. (C.12) and (C.13). Equations (C.18) – (C.22), together with the initialization in Eq. (C.16), constitute the standard Kalman filter recursion for the present model.

Substituting Eq. (C.14) into the chain-rule factorization in Eq. (C.5) gives the exact likelihood for lineage *ℓ*:

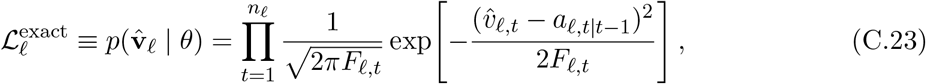

where *a*_*ℓ,t*|*t−*1_ and *F*_*ℓ,t*_ are obtained recursively from Eqs. (C.16)–(C.22).

Assuming that different lineages are conditionally independent given *θ*, the total likelihood is

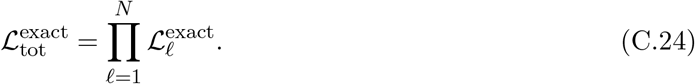

For our model, we can exactly calculate the likelihood from the Kalman filter as the model is a linear Gaussian state-space model: the latent AR(1) dynamics are linear in *v*_*ℓ,t*_, the observation model is linear, and both noise terms are Gaussian. Under these conditions, all predictive and filtered distributions remain Gaussian and are therefore fully characterized by their means and variances.

### C.3 Prior distributions and parameter bounds

To model lineage-to-lineage variability in mean birth size, we assign a hierarchical Gaussian prior to the lineage-specific means. Conditional on the population mean *µ*_*ϵ*_ and the between-lineage standard deviation *σ*_*ϵ*_, the lineage-specific means are independent:

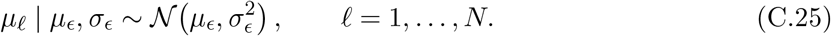

Their joint conditional density is therefore

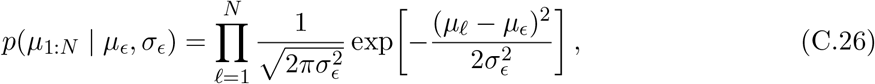

where *µ*_1:*N*_ = (*µ*_1_, …, *µ*_*N*_ ).

We assign mutually independent uniform priors to the remaining parameters:

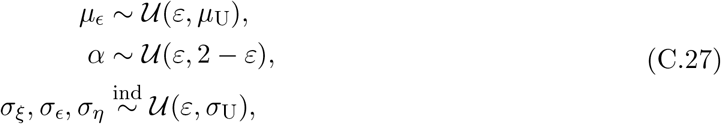

where *σ*_*ξ*_, *σ*_*ϵ*_, and *σ*_*η*_ denote the intrinsic, extrinsic, and measurement-noise amplitudes, respectively. We set *ε* = 10^*−*9^ to exclude zero-valued scale parameters and to restrict *α* to the stationary interval 0 < *α* < 2. The upper bounds are determined separately for each dataset from the pooled observed birth sizes. Specifically, *µ*_U_ is set to ten times the pooled mean birth size across all lineages and generations, whereas *σ*_U_ is set to ten times the corresponding pooled sample standard deviation. These data-dependent bounds are intended to provide broad but finite support for the prior distributions. We verified that the posterior estimates were insensitive to reasonable changes in these bounds.

### C.4 Posterior distribution

Let the full dataset be 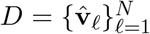 and parameters *θ* = *µ*_1_, …, *µ*_*N*_, *α, µ*_*ϵ*_, *σ*_*ϵ*_, *σ*_*ξ*_, *σ*_*η*_ . Bayes’ rule gives

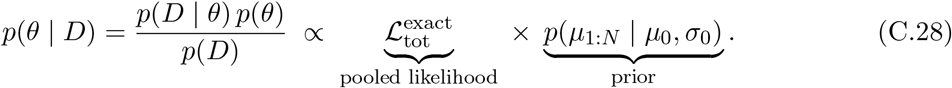

So, the full posterior distribution can be obtained by substituting the likelihood from Eq. (C.24) and prior from Eq. (C.26). The evidence *p*(*D*) is a normalization constant that we do not need to compute if we use MCMC.

### C.5 Inference using Markov chain Monte Carlo (MCMC)

We generate a Markov chain *θ*^(1)^, *θ*^(2)^, … whose stationary distribution is *p*(*θ* | *D*) given in Eq. (C.28). At iteration *k*:

1. Propose a random-walk move *θ*^*′*^ = *θ*^(*k*)^ + Δ ⊙ *ε*, with *ε* ∼ N(0, *I*).
2. Compute the acceptance probability

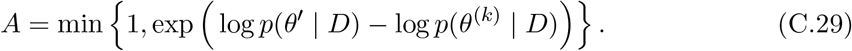
3. Accept *θ*^(*k*+1)^ = *θ*^*′*^ with probability *A*; otherwise set *θ*^(*k*+1)^ = *θ*^(*k*)^.
After burn-in, the retained samples approximate posterior draws. Hence, posterior means are estimated by sample averages, e.g.

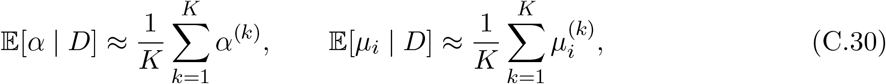

and credible intervals come from posterior quantiles of the samples.

### C.6 Bayes factor for testing a point hypothesis on *α*

We want to quantify the evidence for a prescribed value *α*_0_ against the alternative that *α* is free to vary between [0, 2]. To this end, we compared a point-null model with a more general model using a Bayes factor.

Let *D* denote the observed data. We write

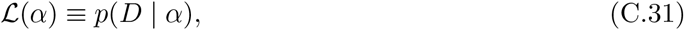

where L(*α*) denotes the likelihood as a function of *α*. If additional parameters are present, then L(*α*) should be understood as the marginal likelihood after integrating out those nuisance parameters.

#### Point-null model

Under the null model *M*_1_, the parameter is fixed at the value *α*_0_. This is represented by the prior

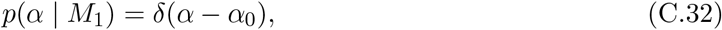

where *δ*(·) is the Dirac delta function. The corresponding model evidence is

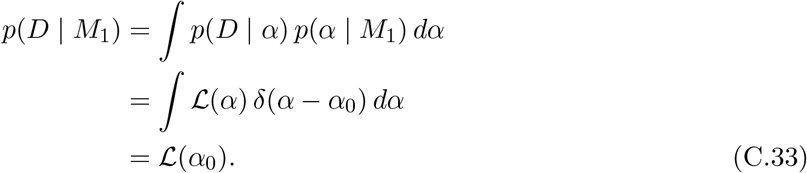

#### General model

Under the alternative model *M*_2_, the parameter *α* is unknown and assigned a uniform prior on [0, 2],

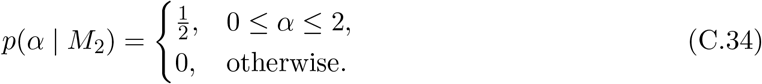

The corresponding evidence is

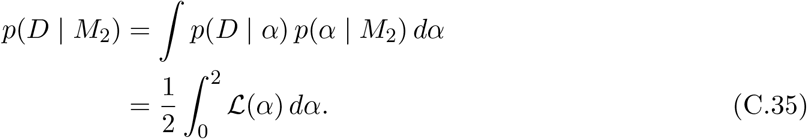

**Bayes factor** The Bayes factor in favor of the point-null model over the general model is

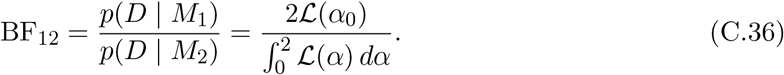

The Bayes factor, BF_12_ ≥ 3.2 is interpreted as substantial evidence for the point hypothesis *α* = *α*_0_.

#### Computation using the Savage–Dickey density ratio

Direct evaluation of Eq. (C.36) requires the marginal likelihood *p*(*D* | *α, M*_2_), obtained by integrating the likelihood in Eq. (C.24) over the lineage-specific means *µ*_1:*N*_, the population mean *µ*_*ϵ*_, and the noise amplitudes *σ*_*ϵ*_, *σ*_*ξ*_, and *σ*_*η*_. Rather than evaluating this high-dimensional integral directly, we use Markov chain Monte Carlo (MCMC) to sample the joint posterior under the general model *M*_2_. Because the point-null model *M*_1_, in which *α* = *α*_0_, is nested within the general model *M*_2_, the Bayes factor can be evaluated using the Savage–Dickey density ratio [46, 47]:

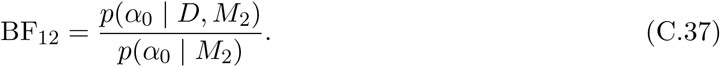

Here, the numerator and denominator are the posterior and prior densities, respectively, evaluated at *α*_0_. Under the general model, *α* ∼ *U*(0, 2), and hence

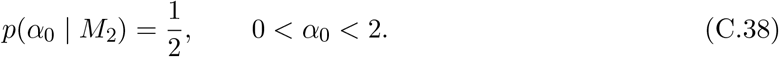

The Bayes factor is therefore

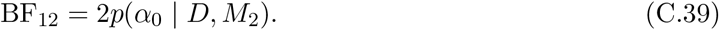

We estimate the posterior density *p*(*α*_0_ | *D, M*_2_) at *α*_0_ from the MCMC samples using kernel density estimation (KDE). KDE is used only to estimate the posterior density required by the Savage–Dickey ratio and is not part of the underlying statistical model.

**Figure C.1:**
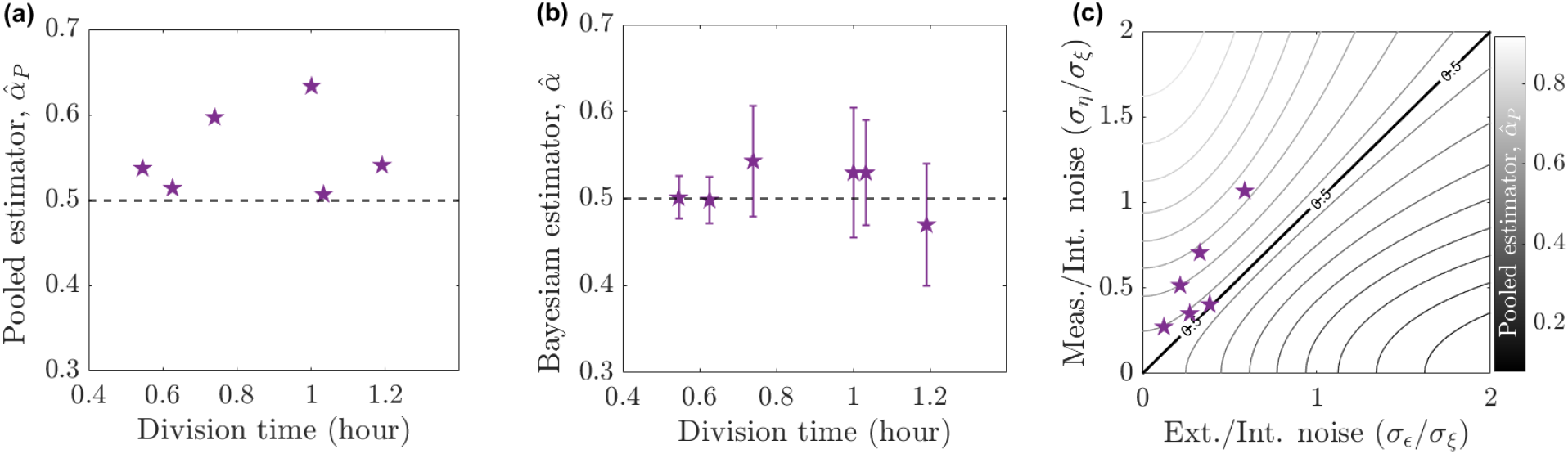
Bayesian inference shows adder cell-size regulation for *B. subtilis* [14]. **(a)** Conventional pooled estimates of the size-regulation parameter, 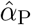, are plotted with mean division time and slightly deviate from adder (*α* = 0.5). (**b**) Corresponding Bayesian estimates of *α*; symbols denote posterior means, and error bars denote 95% credible intervals. Horizontal dashed lines in the estimator panels mark the adder value, *α* = 0.5. (**c**) Each dataset is positioned according to its inferred extrinsic-to-intrinsic noise ratio, *σ*_*ϵ*_*/σ*_*ξ*_, and measurement-to-intrinsic noise ratio, *σ*_*η*_*/σ*_*ξ*_. The shading and contour lines show the analytical expectation of the pooled estimator from Eq. (9) in the main text, assuming an underlying adder with *α* = 0.5. The thick black contour marks 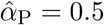, along which the opposing biases from extrinsic variability and measurement noise cancel. Above this contour, measurement noise dominates, biasing 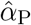 upward; below it, extrinsic variability dominates, biasing 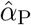 downward. For each dataset, the value of 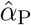 predicted by its location in the noise-ratio plane closely agrees with the observed value in panel (**a**). The differences in the relative amplitudes of measurement and extrinsic noise account for the deviation of the pooled estimator from the adder value.

## D AR(1) dynamics of daughter-cell size in budding yeast

Budding yeast undergoes asymmetric cell division, producing a larger mother cell and a smaller daughter cell [4, 27]. Fig. D.1 illustrates the asymmetric budding-yeast cell cycle and the notation used below. We consider a daughter-lineage in which *v*_*t*_ denotes the birth volume of the daughter cell in generation *t*. The cell grows through the unbudded and budded phases until it reaches the total division volume, 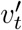 . Division separates the bud from the mother, where the bud becomes the daughter cell of the next generation, with volume *v*_*t*+1_. Although both mother and daughter cells can initiate subsequent budding cycles, we follow only the daughter lineage. If 1*/q* denotes the mean fraction of the division volume inherited by the daughter, then

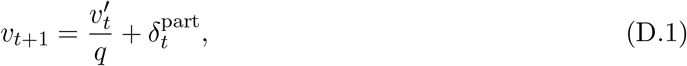

where 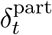 represents partitioning noise. Symmetric division corresponds to *q* = 2, whereas *q >* 2 describes asymmetric division in which the daughter inherits less than half of the division volume. The stochastic size-control relation between the division and birth volume of generation *t* can be written as

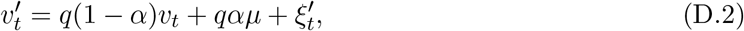

where *µ* is the mean daughter-cell birth volume, *α* characterizes the size-regulation parameter, and 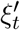 represents fluctuations accumulated before division. Combining Eqs. (D.2) and (D.1) gives the daughter-lineage recurrence

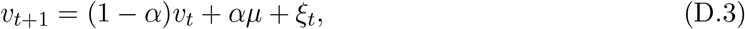

with 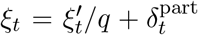. For an adder mechanism, cells accumulate a constant volume from birth to division, leading to a unit regression slope between birth and division volume. Therefore, from Eq. (D.2) we can write

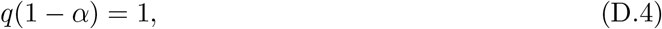

which gives

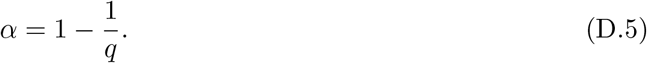

Thus, symmetric division with *q* = 2 gives *α* = 0.5 whereas taking *q* = 3 as a representative asymmetric-budding case gives *α* = 0.67.

**Figure D.1.**
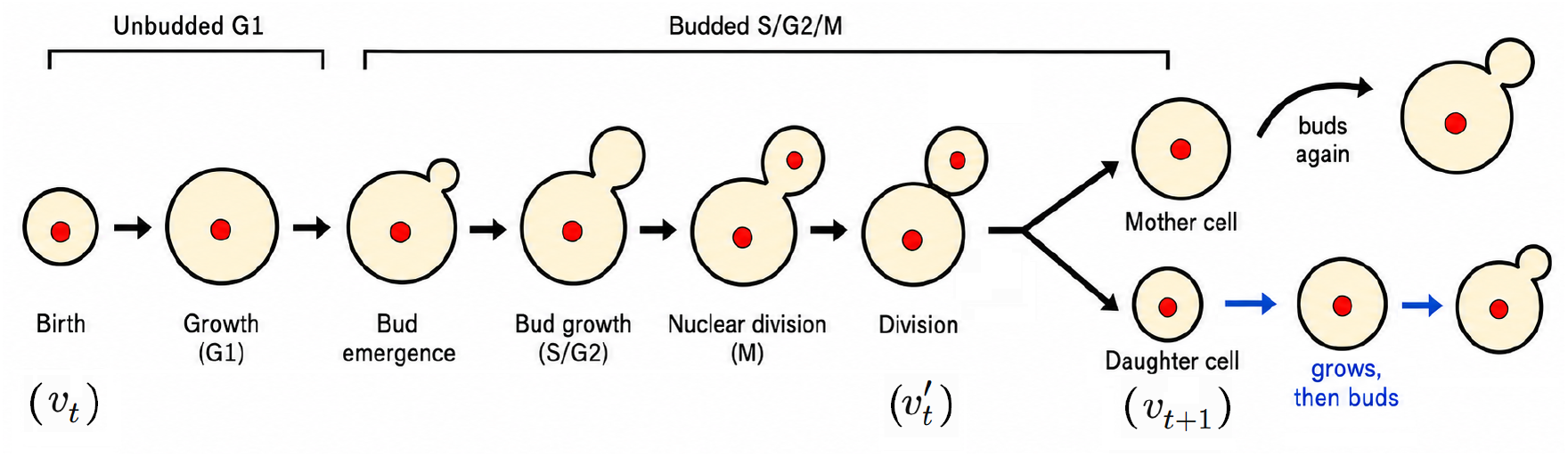
Schematic of the cell cycle of budding yeast. A daughter cell born in generation *t* has volume *v*_*t*_ and grows to the division volume 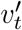. Division produces a mother and a daughter. Both mother and daughter cells can initiate subsequent budding cycles, but we follow only the daughter lineage.

## Notes

### Competing Interest Statement

The authors have declared no competing interest.

